# Concerted evolution and unorthodox recombination of human subtelomeres

**DOI:** 10.64898/2026.07.10.737660

**Authors:** Andrea Guarracino, Angela Gyamfi, Human Pangenome Reference Consortium, Erik Garrison

## Abstract

Human subtelomeres contain duplicated sequence that is shared among the ends of non-homologous chromosomes and provides a substrate for ectopic exchange [1–6]. However, incomplete reference assemblies and chromosome-by-chromosome analyses have prevented a population-scale view of the extent and organization of subtelomeric exchange [7–9]. Here we apply a reference-free pangenome approach to 465 near-complete human assemblies, comparing every chromosome end against every other, and find that high-identity pseudo-homolog regions [10] occur on 41 of 48 chromosome arms. These regions group into sequence communities, sets of chromosome ends that share duplicated sequence. Across these communities, copy-number enrichment is concentrated in gene modules associated with chromosome dynamics, intracellular organization and germ-cell development. Human chromosome-contact maps show preferential proximity between subtelomeres with similar sequences, persisting even in adjacent flanks that lack the shared sequence used to define each pair. Mouse contact maps show the same preference, strongest in meiosis at the zygotene bouquet, when telomeres cluster at the nuclear envelope. In a three-generation telomere-to-telomere pedigree, whole-genome comparison identifies putative recombination between subtelomeric regions on non-homologous chromosomes from the same sequence communities, while recovering the obligate X-Y crossover in PAR1, the pseudoautosomal region shared by the X and Y chromosomes, during male meiosis. These results generalize known subtelomeric exchange systems into a near-ubiquitous architecture and support recurrent ectopic exchange as a genome-wide force in the concerted evolution of human chromosome ends.

---

Human subtelomeres are among the most rearrangement-prone and fast-evolving regions of the human genome [1]. Cytogenetic and clone-based studies established that each chromosome end is a mosaic of duplicated segments also found at the ends of several other, non-homologous chromosomes [2, 3]. These mosaics are organized into two domains: a proximal region of chromosome-specific sequence and a distal zone of repeats shared across many ends. These duplicated blocks harbor subtelomeric copies of gene families and repeat-associated loci (olfactory receptors, tubulins and the D4Z4/DUX4 macrosatellite), with copy number and sequence varying within and between human populations [1, 11, 12]. These chromosome ends also exchange sequence ectopically, between non-allelic sites (often on non-homologous chromosomes), by translocation and gene conversion [4–6], and that exchange has continued into recent human evolution, leaving much of subtelomeric sequence derived from duplications specific to the human lineage [3].

This ectopic exchange has well-known consequences at particular chromosome ends. The pseudoautosomal regions PAR1 and PAR2 are shared by the sex chromosomes, and PAR1 carries the obligate male sex-chromosome crossover [13, 14]. The five acrocentric chromosomes (13, 14, 15, 21, 22), whose short arms carry ribosomal DNA, cluster around the nucleoli, where they recombine and underlie recurrent Robertsonian translocations [10, 15–17]. The 4q35 D4Z4 macrosatellite, homologous to the 10q26 array, is the causal locus in facioscapulohumeral dystrophy, and the DUX4 gene it contains, normally silenced, is reactivated in diverse cancers [18, 19]. Unbalanced subtelomeric rearrangements are also a recurrent cause of intellectual disability [1]. Yet this picture was built from heterogeneous, locus-focused evidence—fluorescence in situ hybridization, BAC and half-YAC clones, monochromosomal hybrids, and complete-assembly studies of particular chromosome classes or genomes [10, 16, 20–22]—rather than a uniform map of exchange across all chromosome ends in a large haplotype-resolved cohort.

Two advances now make this view possible. The telomere-to-telomere CHM13 assembly resolved one human genome end to end [7]. The second release of the Human Pangenome Reference Consortium (HPRC v2) provides near-complete, haplotype-resolved assemblies for 232 individuals from five superpopulations [9]. Our analyses of heterologous acrocentric chromosomes and complete assemblies of human Robertsonian chromosomes established that their short arms recombine through shared pseudo-homolog regions (PHRs) [10, 16]. We extended this reference-free, all-to-all pangenome graph approach to the full cohort and all chromosome ends to map inter-chromosomal sub-telomeric sharing without chromosomal partitioning, measure how that sharing is organized into sequence communities, and test whether those communities coincide with three-dimensional nuclear proximity and ongoing pedigree-scale recombination.

## A whole-genome survey reveals subtelomeric sharing

We began with an unbiased, genome-wide survey of inter-chromosomal sequence sharing. We aligned 25,272 sampled ordered pairs (11.6% of all possible) from a 466-sequence whole-genome set comprising 464 HPRC v2 haplotypes plus CHM13v2.0 and GRCh38. The sample lies well above the modeled connectivity threshold for the assembly-pair network (Methods), allowing sequence sharing to be recovered through transitive paths even when two assemblies were not directly compared. We treated these sampled alignments as an implicit pangenome graph: for each non-overlapping 100 kb CHM13 window we queried the indexed alignments and followed up to five alignment hops, so a match could be reached through an intermediate assembly (A*→*B*→*C) [23]. The resulting genome-wide map reveals high-identity inter-chromosomal sharing concentrated at chromosome ends (Fig. 1A). The acrocentric short arms and pseudoautosomal regions appear as expected positive controls, but the same pattern recurs at most other chromosome ends.

**Fig. 1:**
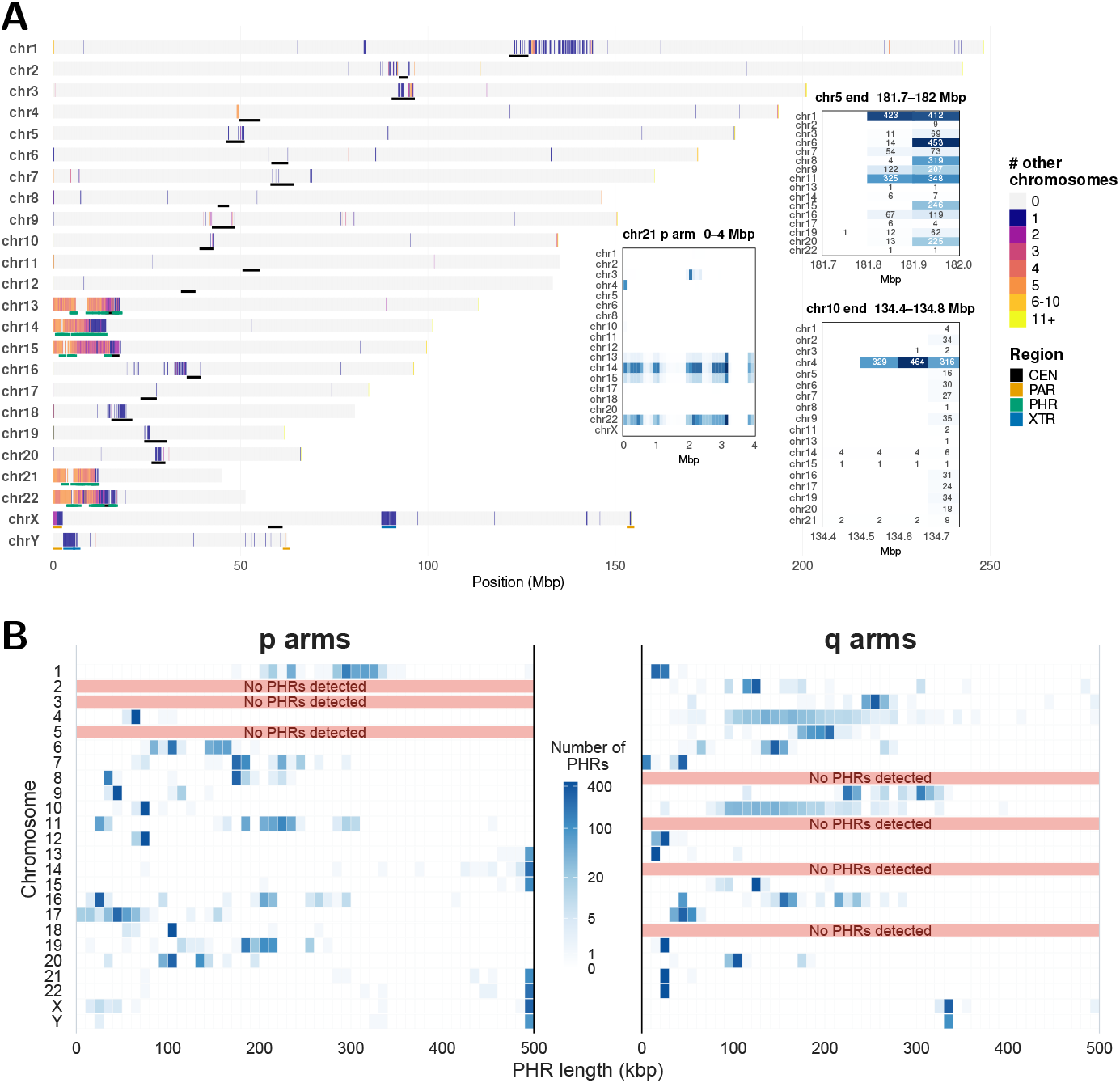
Genome-wide subtelomeric sharing and PHR extent. **(A)** Inter-chromosomal sequence sharing in a whole-genome alignment set comprising 464 HPRC v2 haplotypes, CHM13 and GRCh38. Each CHM13 chromosome is drawn as a track of 100 kb windows colored by the number of distinct partner chromosomes reached elsewhere in the pangenome (0 to 11+), together with a region track (centromere, PAR, PHR [pseudo-homolog region], XTR [X-transposed region]); insets give per-partner haplotype counts at three representative telomere-proximal loci: the chr5 q-end (which shares with chr6 and chr1), the chr21 acrocentric p-arm, and the chr10 q-end (the 4q/10q DUX4 pair). **(B)** Distribution of PHR lengths, shown separately for p-arms and q-arms and colored by the number of PHRs per 10 kb length bin (pseudo-log scale); each PHR is a telomere-anchored interval, so its length is how far the shared sequence extends inward. Saturation at 500 kb reflects the input flank size.

Applying this approach to 465 near-complete assemblies, we identified 15,668 PHRs on 41 of the 48 chromosome arms; the remaining seven arms (chr2p, chr3p, chr5p, chr8q, chr11q, chr14q, chr18q) contained no PHR under our detection criteria. We extracted the 500 kb flank inward from each telomere and aligned the flank set in a separate, complete all-against-all comparison (not the sampled whole-genome survey set), without restricting comparisons to homologous chromosomes, and called a PHR where consecutive windows each contained at least five alignments from each of at least two chromosomes, including the source chromosome, at *≥* 95% identity (Methods). Walking inward from each telomere until inter-chromosomal similarity disappears, we found that PHRs span tens to hundreds of kilobases (median 105 kb, mean 144 kb; Fig. 1B). Excluding the acrocentric short arms and the pseudoautosomal regions, these PHRs total 3.51 Mb of subtelomeric sequence, with the median PHR about a third the length of PAR2 [14].

### Community structure of subtelomeric sharing

To measure how much sequence chromosome ends share, we built an explicit pangenome graph from all 15,668 PHR sequences [24, 25]. Unlike the implicit alignment graph used for the survey, which follows sampled alignments on demand, this graph materializes the aligned sequence structure: shared sequence is represented by graph nodes and each PHR by a path through those nodes. Its main connected component contains 15,654 of the sequences (99.9%) and is too tangled to interpret node by node. We therefore measured pangenome similarity between chromosome ends as the Jaccard overlap of the graph nodes each traverses, and reduced the result to an arm-level similarity matrix (Methods). The matrix is structured: locally dense blocks of high similarity are embedded in a broader continuum of lower inter-community similarity (Fig. 2B, left). Leiden community detection [26], which groups a similarity network into clusters with stronger within-cluster than between-cluster connectivity, partitions the 41 PHR-bearing arms into 15 arm-level communities (Fig. 2B, right). The communities recover every previously described case of inter-chromosomal subtelomere homology: the two pseudoautosomal pairs Xp/Yp and Xq/Yq [27], the five acrocentric short arms [15], the 10p/18p pair first reported by Linardopoulou and colleagues [3], and the 4q/10q D4Z4 pair [18]. Average-linkage hierarchical clustering (UPGMA) provides an alternative display and method comparison (Fig. 2B, left), with exact agreement on 12 of the 15 communities; we do not interpret the ordering as an evolutionary tree. One community, C6, groups six q-arms (22q, 21q, 19q, 1q, 13q, 17q) into a broad enriched neighborhood rather than a set of discrete pairwise relationships (Methods).

**Fig. 2:**
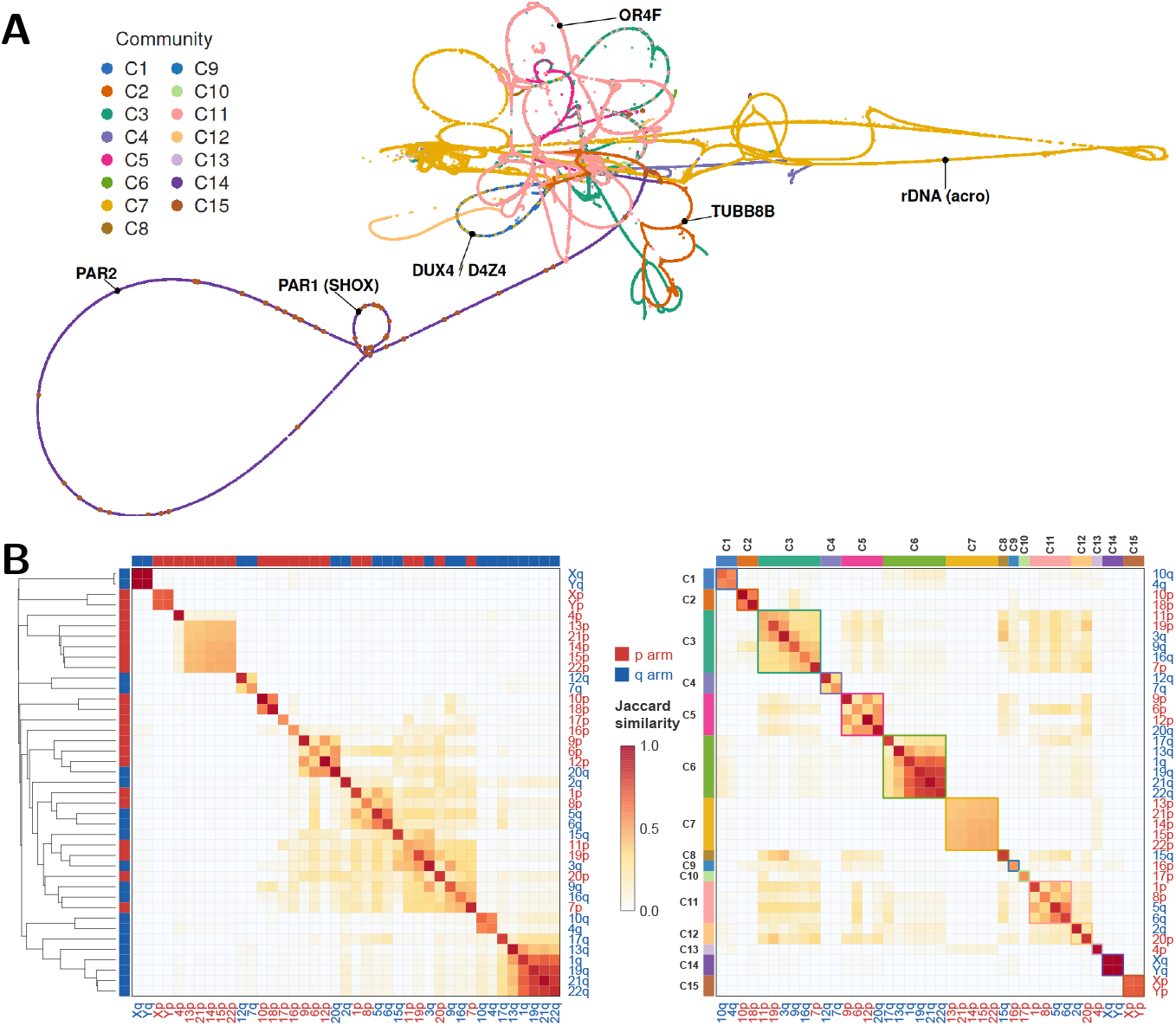
Pangenome similarity structure and communities. **(A)** 2D layout of the all-subtelomere PHR (pseudo-homolog region) graph (main connected component), each node colored by the arm-level community of the PHR paths crossing it (same community palette as panel B); the communities occupy distinct regions of the layout. **(B)** Arm-level Jaccard similarity heatmap (41 *×* 41), shown twice. *Left*, ordered by an average-linkage dendrogram that places arms with similar subtelomeric PHR profiles adjacent (p-arms red, q-arms blue), showing locally dense blocks on a broader similarity continuum. *Right*, the same matrix ordered by the 15 arm-level communities (**Methods**), with community color bands.

At the sequence level, the representative communities shown here are characterized by duplicons, blocks of sequence that recur at multiple chromosome ends and may contain genes, pseudogenes or non-coding sequence (Fig. 3). Community C1 (4q/10q) carries the D4Z4 macrosatellite and its DUX4 retrogene, with homologous DUX4L arrays and FRG1/FRG2 paralogs on both ends and copy number ranging from 0 to 22 across haplotypes (Fig. 3A) [18, 28]. Community C2 (10p/18p) is anchored by a TUBB8-family duplicon, with TUBB8 on 10p and its TUBB8B paralog on 18p amid tandem tubulin-related copies (Fig. 3B) [3]. Community C11 (5q/6q) is built on the OR4F olfactory-receptor family and its pseudogenes (Fig. 3C). More broadly, characteristic subtelomeric duplicons, rather than community-specific protein-coding genes, define the communities [22], and subtelomeric gene content is correspondingly dominated by pseudogenes and non-coding RNAs. Within-community arm pairs are overwhelmingly assortative by arm type: 41 of 58 pairs (71%) join two p-arms or two q-arms, compared with 400 of 820 (49%) among all possible pairs of the 41 PHR-bearing arms. PHR sharing is therefore constrained by chromosome-arm context rather than distributed indiscriminately among chromosome ends.

**Fig. 3:**
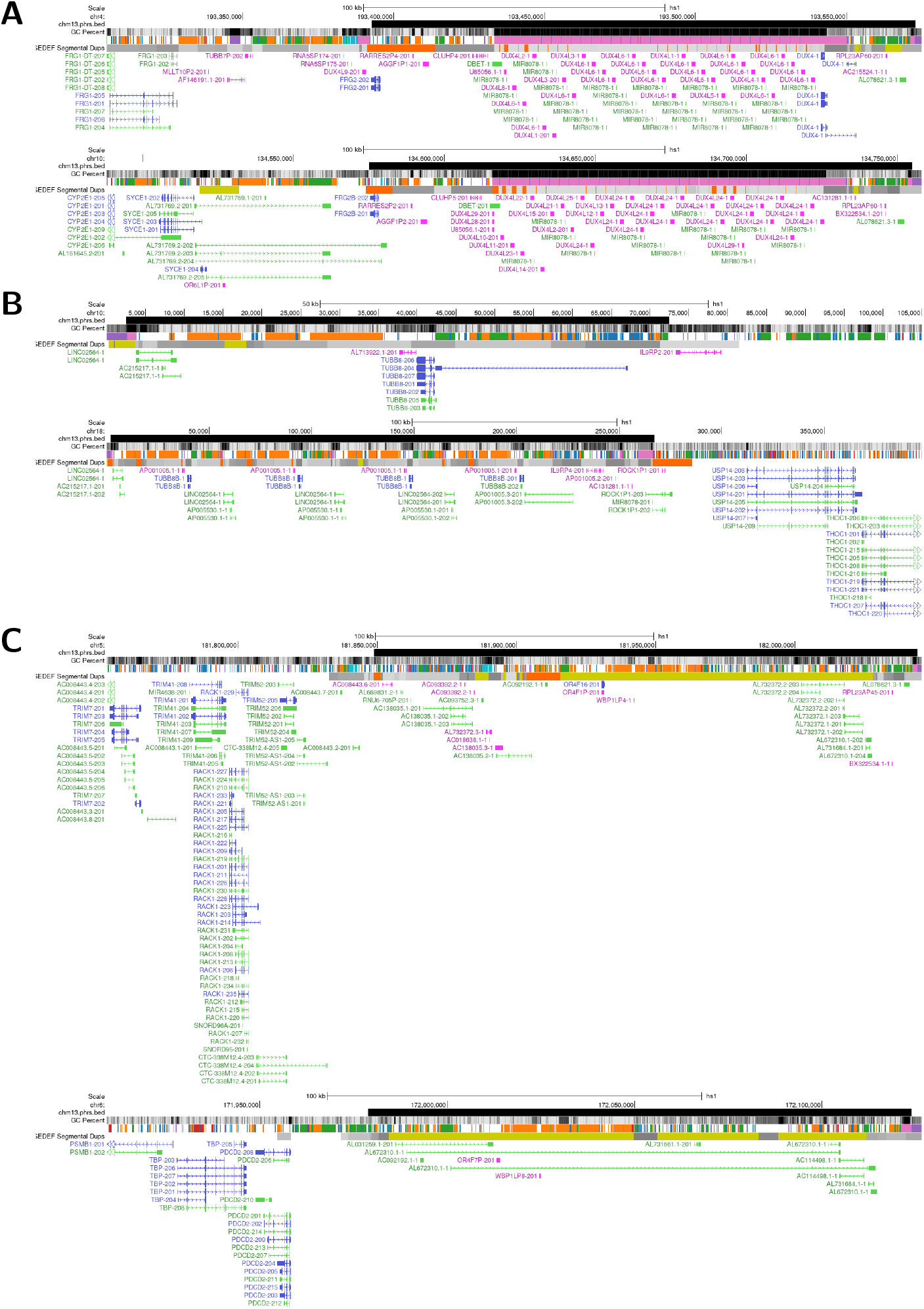
Architecture of representative communities. Sequence-level views (CHM13 coordinates, with PHR intervals, GC%, segmental duplications and gene models) of three communities, so the duplicon content is legible without a genome browser. **(A)** Community C1 (4q/10q), the D4Z4 macrosatellite and DUX4 retrogene: homologous DUX4L arrays and FRG1/FRG2 paralog families on the 4q35 (top) and 10q26 (bottom) ends, copy number 0–22 across haplotypes. **(B)** Community C2 (10p/18p), a TUBB8-family block: TUBB8 on 10p (top) and its TUBB8B paralog amid tandem tubulin-related duplicons on 18p (bottom). **(C)** Community C11 (5q/6q; 5q top, 6q bottom), built on the OR4F olfactory-receptor family and its pseudogenes.

### Functional enrichment within subtelomeric PHRs

We performed a copy-number-aware enrichment analysis of Gene Ontology (GO) [29] and Reactome [30] annotations in CHM13 to assess the functional significance of the PHR compartment (Methods). Gene copies within PHRs were compared with all annotated gene copies elsewhere in the genome, with each duplicated copy counted separately rather than collapsed by gene name or family. The analysis identified 209 significantly enriched terms spanning biological processes, molecular functions, cellular components and Reactome pathways.

The enrichments fall into several functional systems whose distributions differ markedly across the PHR communities (Extended Data Fig. ED2). The most community-restricted system is associated with DUX4: functions related to zygotic genome activation, transcriptional regulation, the nuclear envelope and the cell cycle are concentrated in the community C1 exchange network linking 4q and 10q. By contrast, functions involved in intracellular organization and chromosome dynamics are distributed across multiple communities. A recurrent WASH/DDX11 module spans six communities, combining WASH-associated endosomal actin and exocyst activity with DDX11-associated helicase activity and sister-chromatid cohesion. A second distributed system is associated with male germ-cell development and sexual reproduction. The direct GO term spermatid development is enriched 25-fold relative to the rest of the genome and is carried by 19 SEPTIN14-associated PHR copies; broader enriched terms include spermatogenesis, germ-cell development, gamete generation and sexual reproduction. These SEPTIN14-associated copies also contribute septin and cytokinesis functions and are concentrated in community C3, with additional copies in communities C5, C11 and C12. Signaling functions associated with WBP1L/CXCL12 and ribosomal and nucleolar functions associated with RPL23A are also shared among several otherwise distinct communities.

Extended Data Fig. ED2 maps these systems across chromosome ends and shows that PHRs do not carry a uniform functional repertoire. Instead, each community contains a different combination of partially overlapping modules: some are highly amplified within a single exchange network, whereas others recur across multiple communities. This functional mosaic parallels the underlying sequence-sharing communities and suggests that inter-chromosomal exchange redistributes discrete blocks of gene content rather than homogenizing entire chromosome ends.

### Subtelomere similarity mirrors physical proximity

To explore the three-dimensional configuration of these regions, we measured inter-chromosomal contact at PHR coordinates in Hi-C, Pore-C and CiFi maps. These assays quantify contact frequency between genomic regions as a proxy for nuclear proximity. Across all of them, contact between subtelomeres increases with their PHR sequence similarity. We measured this per inter-chromosomal PHR pair within a single genome in HG002 Pore-C (Fig. 4A; pointwise Spearman *ρ* = 0.376, *n* = 2,830), with contact increasing similarly in CHM13 Hi-C, HG002 Hi-C and HG002 CiFi (Extended Data Fig. ED1). Because near-identical PHRs on non-homologous chromosomes cannot be told apart by read mapping, we interpret contact at a PHR window as an aggregate over its near-identical copies rather than reads uniquely assigned to a single PHR. The 100 kb immediately centromere-ward of each PHR, the non-PHR flanks, consists of unique sequence and so provides the key control for this multi-mapping concern. In HG002 Hi-C, the ratio of between-to within-community contact (*B/W*; lower is stronger) is 0.0013 for the flanks versus 0.028 in the PHR windows, a 21-fold stronger within-community enrichment, although the flanks do not contain the shared sequence used to define PHR pairs. A Pore-C contact matrix [31] ordered by sequence community shows the same community block structure after observed/expected inter-chromosomal normalization, which accounts for baseline differences in inter-chromosomal contact so that broad nuclear-contact biases do not masquerade as community structure (Fig. 4B) [32, 33]. The same within-community enrichment holds in the HG002 CiFi contact map [34]. Subtelomeric sequence similarity is positively correlated with contact across all four stages of prophase I in mouse spermatocytes (pointwise Spearman *ρ* = 0.419, 0.614, 0.576 and 0.245 at leptotene, zygotene, pachytene and diplotene; all *n* = 1,135; Fig. 4C), strongest at the zygotene bouquet—when telomeres transiently cluster at the nuclear envelope—and lowest at diplotene, once the bouquet disperses. We compared stage-resolved Hi-C from inbred C57BL/6 (B6) spermatocytes mapped to a combined B6 and CAST T2T reference [35, 36], holding assemblies, PHR sequence identities and read-placement procedure constant across stages. In mouse, where detectable PHRs are concentrated on p-arms, the zygotene peak is likewise a p-arm feature. Restricting to inter-chromosomal pairs of the same arm type, the correlation peaks at zygotene over pairs of two p-arm PHRs (Spearman *ρ* = 0.285, 0.508, 0.323, 0.023; *n* = 722), whereas over pairs of two q-arm PHRs it peaks at diplotene (*ρ* = 0.352, 0.338, 0.406, 0.420; *n* = 42)—though the q-arm subset is small and p-arm windows carry broader arm-context effects, so we do not read the arm split as a clean arm-type experiment.

**Fig. 4:**
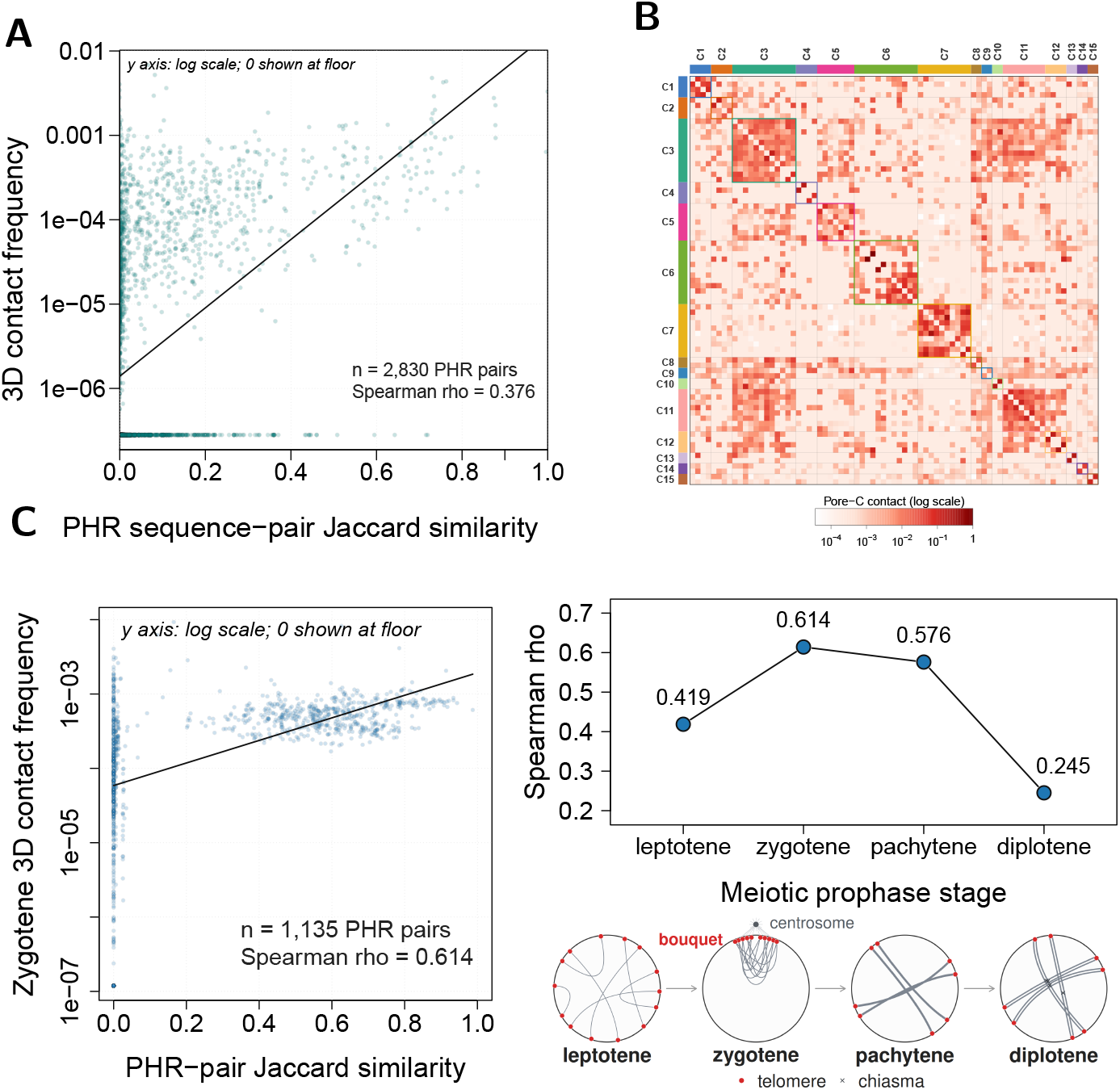
Sequence similarity tracks three-dimensional proximity. **(A)** Each dot is one inter-chromosomal pair of HG002 pseudo-homolog regions (PHRs); *x*, their Jaccard sequence similarity in the pangenome graph; *y*, their length-normalized 3D contact at the PHRs’ exact coordinates in the same HG002 Pore-C map at 20 kb (log scale, zero-contact pairs at the axis floor; line, linear fit). **(B)** HG002 Pore-C contact matrix at 20 kb, ordered by sequence community after observed/expected inter-chromosomal normalization; within-community contacts are enriched (*B/W* = 0.054, between/within). **(C)** Mouse meiotic Hi-C (PHRs established de novo from B6 and CAST T2T assemblies; 20 kb contacts over 1 Mb flanks). Left, one dot per inter-chromosomal mouse PHR sequence pair, with zygotene contact (per-bin-pair length-normalized as in (A), log) versus PHR Jaccard similarity. Right, the per-PHR-pair Spearman statistic across leptotene, zygotene, pachytene and diplotene.

### Ectopic subtelomeric recombination within a pedigree

We traced transmission through the WashU pedigree [37], the only one with complete telomere-to-telomere assemblies in all three generations (Fig. 5A), to detect recent exchange in a single family. We compared the chromosomes a daughter inherited from her father against both of his haplotypes by whole-genome alignment (Methods, Fig. 5). This father-to-daughter transmission is a useful positive-control setting because recombination is biased toward chromosome ends in the male germline [38], and the same procedure recovers PAR1 sharing between Xp and Yp, the obligate site of male sex-chromosome recombination. Almost everywhere, each of the daughter’s chromosomes matches its homolog in her father. However, the whole-genome scan (Fig. 5C) highlights two putative autosomal PHR exchanges, chr5q/chr1p and chr9q/chr3q, where a subtelomere’s best match is instead a non-homologous chromosome in her father. For chr9q/chr3q, the best match shifts from the father’s chr9 to his chr3q across the 20 kb tract (Fig. 5B). The two autosomal segments are short, 20–28 kb, compared with 138 kb for the PAR1 control. The exchanged tracts are tens of kilobases. This is well below the scale of whole-arm translocations and within the range of individual PHRs and their functional modules, consistent with block-scale rather than wholesale exchange. Each exchange pair falls into a community observed in the pangenome survey (Xp/Yp in C15, chr5q/chr1p in C11, chr9q/chr3q in C3). We applied the same scan to two further transmissions: the chromosomes the daughter inherited from her mother, and the chromosomes the granddaughter inherited from that same daughter. The daughter’s maternal side shows only the expected acrocentric-acrocentric sharing (Fig. 5D), while the granddaughter’s is again dominated by acrocentric-acrocentric sharing, with two further highlighted candidates, the subtelomeric chr5q/chr1p and a chr21p/chr4p run whose chr4 partner is pericentromeric rather than subtelomeric (Fig. 5E). Across these transmissions, the strongest non-acrocentric subtelomeric signal occurred in the paternal comparison, consistent with the male-specific concentration of recombination at chromosome ends [38]. Acrocentric-acrocentric sharing appears in both germlines, consistent with the rDNA exchange system [10].

**Fig. 5:**
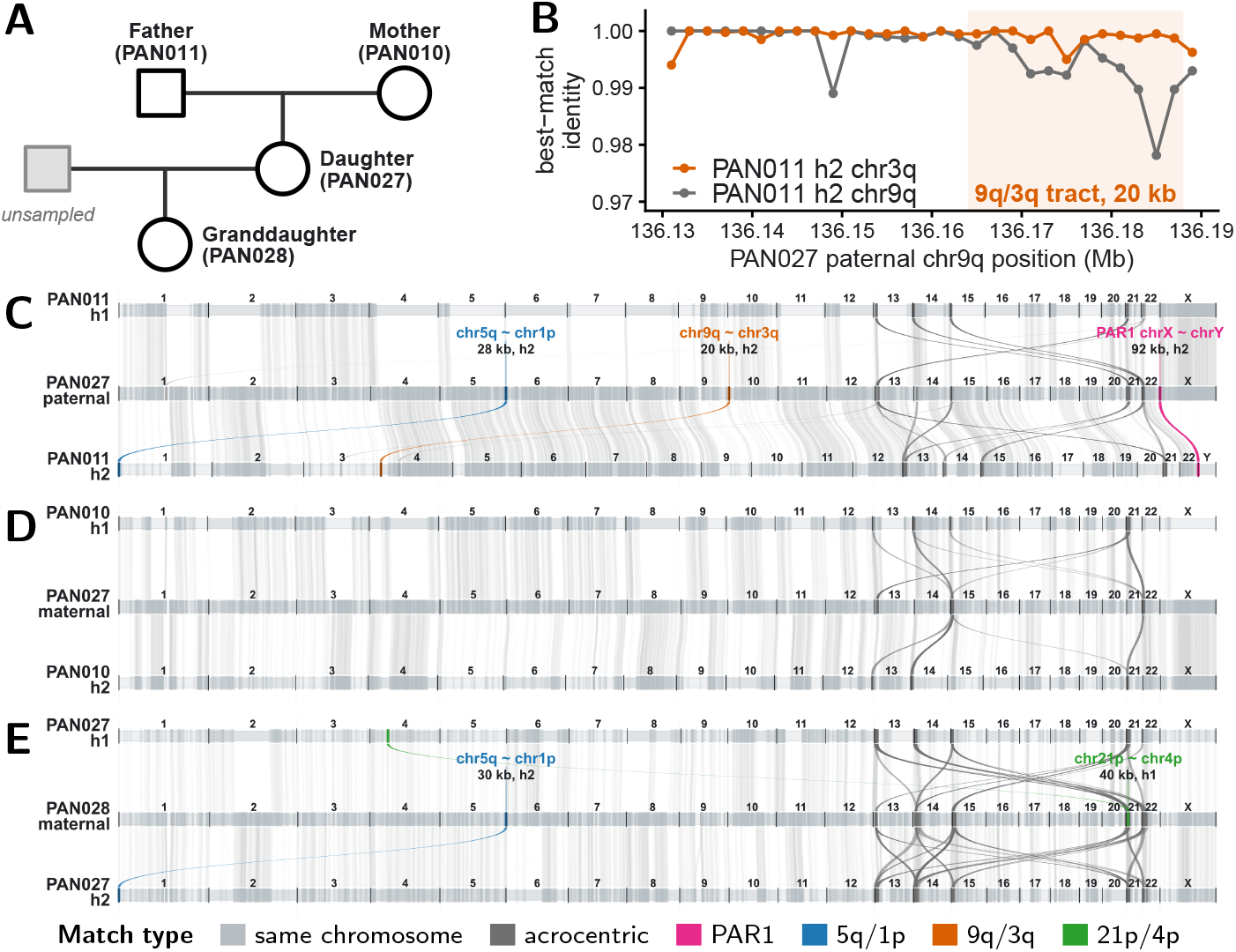
Whole-genome pedigree views of putative subtelomeric exchange. **(A)** The WashU three-generation pedigree, with grandparents PAN011 and PAN010, their daughter PAN027 (the mother of PAN028) and the granddaughter PAN028; PAN028’s unsampled father is drawn as an open square. **(B)** Identity-crossover zoom of the 9q/3q exchange. Along the PAN027 paternal chr9q subtelomere, the bestmatch sequence identity to the father’s own chr9q (gray) drops across the called 20 kb tract (shaded), while the match to the father’s chr3q (orange) stays near-identical. **(C)** PAN027 paternal haplotype compared with the two PAN011 father haplotypes. **(D)** PAN027 maternal haplotype compared with the two PAN010 mother haplotypes. **(E)** PAN028 maternal haplotype compared with the two PAN027 mother haplotypes. In each ribbon panel, chromosomes are concatenated in order and scaled by their length; the middle track is the meiotic product inherited by the child/query haplotype, and the upper and lower tracks are the two parental haplotypes being compared. Light-gray ribbons show same-chromosome homologous chains, while colored ribbons show runs where the best non-homologous parental match beats the same-chromosome or homologous match in the same whole-genome scan (**Methods**).

The two autosomal candidates (chr9q/chr3q and chr5q/chr1p) are consistent with inter-chromosomal exchange. Of the two, chr9q/chr3q is the better-supported call: its chr3q match exceeds the homologous chr9q by a wider identity margin than the chr1p match exceeds chr5q (Methods). In the WashU pedigree crossover map this paternal transmission has no crossover on chromosome 3 [37], so ordinary chr3 recombination cannot account for the chr9q/chr3q signal. We therefore treat these as candidate PHR exchanges consistent with recombination, not as confirmed events with resolved breakpoints.

## Discussion

Together, these results define a subtelomeric exchange compartment in the human pangenome, a distributed set of chromosome ends that share high-identity sequence, sit in preferential three-dimensional proximity, and recurrently exchange material. Individual pieces of this compartment have long been recognized through cytogenetics and locus-specific studies, but had not been assembled into a systematic, population-scale view across nearly complete human genomes. The HPRC v2 assemblies let us build one, turning scattered observations into a sequence-resolved map across 41 chromosome arms, in which the pseudoautosomal, acrocentric, 10p/18p and 4q/10q systems stand out as local peaks within a broader continuum. The 3.51 Mb of subtelomeric sequence mapped here traces the routes along which chromosome-end sequence has repeatedly been copied, exchanged and homogenized [39, 40].

This compartment is also internally organized. Communities assort by arm type, giving pronounced p–p and q–q structure whose cause is not yet clear. But the broader pattern is consistent with a simple, self-reinforcing model: telomere clustering at the meiotic nuclear envelope brings similar ends into contact, and proximity may favor ectopic recombination that helps keep their sequences homogenized. Shared sequence and shared function then feed back to promote co-localization, so that sequence, function and three-dimensional proximity continually reinforce one another. The acrocentric short arms show the loop most cleanly: their shared rDNA draws them to the nucleolus, and nucleolar contact drives the ectopic recombination that homogenizes them [10]. We see the same logic operating on other functional modules outside the NORs. This concerted process operates at a characteristic scale. The exchanged blocks are on the order of tens to hundreds of kilobases, large enough to carry an intact functional module, but small enough to leave chromosome identity intact rather than translocating whole arms. We used copy-number-aware ontology enrichment as a descriptive way to summarize which annotated gene modules are disproportionately represented in the PHR compartment. Because duplicated copies are counted separately, these results describe annotation burden across physical gene copies rather than independent functional systems. The copy-number-aware enrichment pattern reflects this block-scale organization, and the systems that resolve all map onto chromosome inheritance across generations: sister-chromatid cohesion and chromosome organization, germ-cell development, and genome reactivation after fertilization. The dispersal of multiple related copies across chromosome ends could buffer against loss of any one copy, while recurrent exchange could both maintain similarity and generate new variants [3, 22]. PHRs may therefore provide both a stable foundation for chromosome transmission and the flexibility needed for germline systems to accommodate continuing change in chromosome-end sequence and organization.

Which factor initiates the loop—sequence similarity, shared function, or physical proximity—remains unresolved, and the acrocentric example, where rDNA both drives and results from nucleolar clustering, suggests there may be no single starting point. The current evidence has gaps: human contact maps are somatic rather than meiotic, and the meiotic-stage data come only from mouse. Improved evidence may shed light on these questions: germline-stage contact maps and larger multigenerational pedigrees with complete, haplotype-resolved telomere-to-telomere assemblies [37], together with orthogonal assays such as FISH [6]. By applying reference-free pangenome methods to the 465-assembly HPRC v2 panel, we convert relationships inferred from cytogenetics decades ago into a population-scale view of concerted evolution in human subtelomeres.

## Acknowledgments

We thank Heather Mefford for discussion of her work on cytogenetic evidence of subtelomere homology. We thank Rob Williams, Pjotr Prins and Vincenza Colonna for support and discussion. We acknowledge the Human Pangenome Reference Consortium for producing the HPRC v2 assemblies and associated data resources that made this analysis possible. Garrison and Guarracino are funded by NSF PPoSS Award #2118709; NIH R01HG013618; NIH U01HG013760; NIH U01DA057530; NIH U41HG010972; and NIH R01HG013017. Computational resources for this work were supported by the UTHSC Center for Integrative and Translational Genomics.

## Author Contributions

A.Gu. and E.G. conceived and designed the study, developed the computational pipeline, performed the analyses, and interpreted the results. A.Gy. conducted the literature review and performed the functional enrichment analysis. A.Gu., A.Gy., and E.G. wrote the manuscript.

## Competing Interests

The authors declare no competing interests.

## Methods

### Whole-genome implicit pangenome survey

The dataset comprised 464 haplotype-resolved HPRC v2 assemblies from 232 individuals plus CHM13v2.0, for 465 near-complete assemblies, with super-population labels assigned at the HPRC individual level [7, 9]. For the whole-genome survey we added GRCh38, giving 466 input sequences, and used wfmash v0.23.0 [41] (-p 95-P inf) to align 25,272 sampled ordered target– query pairs, representing 11.6% of the 466^2^ possible ordered pairs. We set this sample size using the Erdös–Ŕenyi random-graph model. Under that model the network of 466 sequences is connected with near-certainty (probability above 99.99%) once about 3.5% of the possible pairs are drawn. The 11.6% sampled here is well above this connectivity threshold. Because alignment is symmetric, sampling 11.6% of the ordered pairs connects about 22% of the unordered sequence pairs, which exceeds the model’s diameter-two thresh-old for this graph 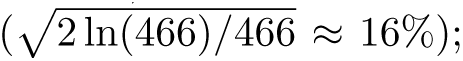 any two sequences are therefore expected to be linked through at most one intermediate. We indexed the resulting pairwise alignments with IMPG v0.4.1. For Fig. 1A, we queried the implicit pangenome graph in non-overlapping 100 kb windows spanning CHM13, following at most five transitive alignment hops at a minimum estimated identity of 98%, and summarized for each window the distinct chromosome origins and mean identity of the sequences reached [23].

### Exclusions and flank extraction

GRCh38 lacks complete subtelomeres and the CHM13v2.0 Y chromosome has its PAR1 masked, leaving neither with telomere-anchored ends, so we excluded both from flank extraction. We detected telomeres with seqtk 1.4-r130 (telo) [42] and classified each contig at least 1 Mb long by the chromosome arm it spans, its p-arm, its q-arm, or both arms when it crosses the centromere. At every telomere-bearing end we trimmed the telomeric repeat and retained the 500 kb immediately internal to it as the flank, so a centromere-spanning contig yields both a p-arm and a q-arm flank, producing 18,827 candidate flanks across the 48 arms. We then excluded the chr18q flank of NA18982 haplotype 1 (contig JBKABS010000018.1) as a scaffolding chimera, since in the whole-genome wfmash alignments its first 83.37 Mb map to chr18 and a terminal 966 kb (83.38–84.35 Mb) map to chrX PAR1, joined across a 100 bp gap of Ns, leaving 18,826 flanks for the all-against-all alignment and PHR calling.

### Alignment, implicit pangenome graph and PHR detection

We aligned all 18,826 flank sequences all-against-all with wfmash v0.23.0 [41] at 95% minimum identity. Every flank was therefore compared with every other without restricting comparisons to homologous chromosomes. We indexed this pairwise alignment set with IMPG v0.4.1 for direct interval queries [23]. Starting at each telomere, we queried 5 kb windows in 5 kb steps, stopping after four consecutive windows failed the detection criteria. A window passed when the direct IMPG query returned at least five alignments from each of at least two chromosomes, including its own (so pseudoautosomal Xp/Yp and Xq/Yq, with a single partner, are also detected), at a minimum identity of 95% and minimum output length of 3 kb. This defined 15,668 PHRs from the 18,826 retained flanks (83.2%).

### Pangenome graph, similarity and community detection

We extracted the 15,668 called PHR intervals and supplied those sequences as the input to pggb v0.7.4 (commit 498c5d7, with wfmash v0.23.0, seqwish v0.7.11, smoothxg v0.8.2, odgi v0.9.2, GFAffix v0.2.1 and vg v1.62.0) at 95% identity, giving an explicit graph with 477,726 nodes and 786,318 edges. We computed all-pairs length-weighted Jaccard similarity from this graph with odgi, giving a 15,668 *×* 15,668 matrix [24, 43–45]. We computed a 2D layout of the main connected component with odgi and rendered it in R (Fig. 2A), each node colored by the dominant arm-level Leiden community of the PHR paths crossing it. The arm-level distance matrix is the mean pairwise Jaccard distance between PHR sequences within each arm pair. From it, arm-level communities were inferred with the Leiden algorithm (R, igraph) after converting distances to graph weights *w_ij_* = exp(*−d_ij_/*median(*d*)) and scanning Leiden resolution from 0.1 to 3.0; we selected the maximum-silhouette partition, 15 communities at resolution 1.16, and the same community count held across resolutions 1.13–1.18. We classified every pair of arms assigned to the same community as either same-type (p–p or q–q) or mixed (p–q) and compared the within-community counts with the corresponding counts across all ^41^ = 820 possible pairs of PHR-bearing arms.

### Copy-number-aware gene ontology enrichment

We counted gene copies from the CHM13v2.0 RefSeq/Liftoff v5.2 annotation (61,312 gene records), treating every genomic copy as a separate unit: copies sharing a gene name or family were kept distinct, so a gene present at 20 positions counted as 20 copies rather than one collapsed family entry. Before looking at any PHR coordinate, we decided which gene’s annotation each copy was allowed to carry: a copy used its own annotation, or, if it had none of its own, inherited terms only through a single explicitly directed NCBI Gene “related functional gene” link; copies with ambiguous, type-only or unsupported evidence were left unannotated and contributed no terms. This kept the number of genomic copies separate from the source used to annotate them. Direct Gene Ontology assignments came from the NCBI human gene2go snapshot of 13 July 2026, propagated through the go-basic release of 15 June 2026; Reactome direct assignments and their parent pathways came from release 96 (1 April 2026), and the NCBI Gene registry snapshots were dated 16 July 2026 [29, 30, 46]. We tested each copy’s directly assigned terms and, separately, the broader parent terms above them in the ontology hierarchy, and fixed the full set of 31,235 database, annotation-level and term combinations before splitting copies into PHR and non-PHR, so that no term was chosen after seeing its result.

A copy entered the analysis if it carried at least one GO or Reactome assignment, giving a fixed universe of *N* = 31,966 copies. We called a gene a PHR gene when its annotation midpoint fell inside a CHM13 PHR, which placed *K* = 187 copies in PHRs and 31,779 outside; a further 215 of the 402 copies with midpoints in PHRs had no confident annotation, and we kept them as a coverage limitation. The rest of the genome was the background, with no sampling, permutation or spatial matching. For each term we compared the number of PHR copies carrying it against the genome-wide count with the exact one-sided hypergeometric upper tail over this fixed population (size *N, K* PHR copies, and the genome-wide count of term-bearing copies), an exact finite-population test of term-bearing physical-copy excess. Because many PHR copies are near-identical paralogs from the same amplified family, they are not independent biological replicates; the *p*-value quantifies evidence for copy-burden excess, not the number of independent functional systems. As the effect size we report the PHR and non-PHR copy fractions, the fold enrichment and odds ratio (both raw and with 0.5 added to each cell, the Haldane–Anscombe correction, so a term absent from the background still gives a finite value), and the observed-minus-expected PHR copy burden.

We controlled multiple testing with Benjamini–Hochberg within each of the four annotation families (GO biological process, molecular function, cellular component, and Reactome) and with Holm across all 31,235 tests, calling a term enriched when its within-family *q ≤* 0.05 and global Holm-adjusted *P ≤* 0.05. We used an any-overlap definition of a PHR gene only to check sensitivity to PHR boundaries. After the tests were fixed, we mapped the contributing copies back to the 15 arm communities and grouped related terms into the six functional classes shown in Extended Data Fig. ED2.

### Hierarchical clustering and resampling controls

For display and method comparison we also ordered the arm-level matrix by average-linkage hierarchical clustering (UPGMA) with hclust (R stats); cutting the dendrogram into 14 clusters gave a mean silhouette of 0.342 (silhouette, R cluster package) and exact agreement with 12 of the 15 Leiden communities, and we use this dendrogram as a heatmap ordering, not as a phylogenetic model. To assess the sharpness of the C6 boundary, we bootstrap-resampled the PHRs and rebuilt the arm-level trees (R package ape). The exact six-arm grouping was unstable under this coarse resampling, consistent with C6 representing a broad, softly bounded neighborhood in the similarity matrix. Because this resampling does not reconstruct the full 15,668-sequence Jaccard matrix, it is not a test of the Leiden partition itself.

### Hi-C, Pore-C and CiFi pipeline

We reprocessed Hi-C, Pore-C and CiFi from raw reads against the corresponding T2T reference (CHM13v2.0 or the matched HPRC v2 haplotype), keeping multi-mapping reads. Standard pipelines apply a mapping-quality filter that drops multi-mapping reads, but subtelomeric PHRs are so similar across chromosomes that their reads map to several places at once, so this filter removes exactly the signal we analyze. We processed Pore-C (NlaIII) and CiFi (DpnII) with pore-c-py 2.1.5, minimap2 2.30 and pairtools 1.1.3, and Hi-C with HiC-Pro 3.0.0 and Bowtie2 2.5.3, binning contacts with cooler 0.10.4 and samtools 1.23 under Python 3.12.12 (numpy 1.26.4, pandas 3.0.0, scipy 1.17.0). Because we apply no mapping-quality filter, a read mapping equally well to several paralogous ends is kept and placed at one of them at random rather than discarded. PHR-internal contacts are therefore aggregate signal over these near-identical ends, not reads confidently assigned to a single end. We generated contact matrices at 5, 10, 20, 50 and 100 kb resolution. For the per-PHR-pair analyses (Fig. 4A, Extended Data Fig. ED1 and Fig. 4C) we summed the balanced contacts over the valid bin-pairs spanning the two PHR windows and divided by the number of valid bin-pairs, giving the mean contact per bin-pair. This normalization removes differences in contact opportunity caused by different numbers of valid bins, so pairs spanning more bins do not automatically have larger total contact.

We report each Spearman correlation as a descriptive measure of how strongly similarity tracks contact, not a p-value, because the pairs are not independent (the same subtelomere recurs in many pairs).

Community-ordered contact matrices used observed/expected inter-chromosomal normalization [32, 33]. For the community-level contact analysis we classified each inter-chromosomal arm pair as within-community or between-community using the arm-level Leiden (*k* = 15) partition, and summarized the contrast as *B/W*, the mean contact of between-community pairs divided by the mean contact of within-community pairs (so *B/W <* 1 indicates stronger within-community contact). As a unique-sequence control for multi-mapping we repeated the calculation on the 100 kb immediately centromere-ward of each PHR, outside the called PHR; these non-PHR flanks show the same within-community enrichment.

### Mouse pipeline

We repeated the entire pipeline on a non-human mammal, using the B6 and CAST T2T assemblies [35]. We extracted telomere-anchored flanks and called PHRs de novo over 1 Mb flanks. The stage-resolved meiotic Hi-C is from inbred C57BL/6 (B6) spermatocytes [36]. Because these mice are inbred and homozygous, the reads are single-haplotype and are not phased into parental haplotypes; we mapped them to the combined B6+CAST T2T reference, keeping multi-mapping reads exactly as in the human pipeline. For each meiotic prophase stage we computed the per-PHR-pair Spearman correlation between length-normalized contact and PHR Jaccard similarity over 1 Mb flanks (*n* = 1,135 inter-chromosomal pairs per stage; Fig. 4C). Each stage gives a single *ρ* value; we did not test whether the differences between stages are statistically significant. We also computed this correlation within each arm class, using only inter-chromosomal pairs in which both PHRs lie on p-arms (*n* = 722) or both on q-arms (*n* = 42).

### Pedigree direct-alignment windows

For each transmission in Fig. 5C–E we aligned one child haplotype (the query) against the combined two-haplotype assembly of the transmitting parent (the target) [37]. Starting from a whole-genome sweepga 0.1.1 [47] alignment (frequency threshold 32, many-to-many mappings, no scaffold jumps), we filtered to 10:10 mapping classes and summarized the query in non-overlapping 2 kb windows, excluding centromeric windows, windows without alignment support, and windows with inter-chromosomal alignment depth above 100. For each retained window we computed impg 0.4.1 similarity and kept the best same-chromosome or homologous parental match and the best non-homologous parental match, assigning the window to the non-homologous class only when the non-homologous match was the most similar. The identity margin of a called exchange is the amount by which its best non-homologous match exceeds the best same-chromosome or homologous match across the tract; a wider margin indicates stronger support. We joined two neighboring windows into one run only when their query and donor intervals were directly adjacent, touching end to end with no gap, on the same donor chromosome, haplotype and sequence; we did not bridge gaps or chain non-adjacent windows. We drew runs of at least 10 kb with mean identity at least 0.95, shown as the non-homologous (colored) and same-chromosome (gray) runs in Fig. 5C–E.

## Data and code availability

HPRC v2 sequencing data and assemblies can be downloaded from https://github.com/human-pangenomics/hprc_intermediate_assembly/tree/main/data_tables. Analysis code, manuscript source, figure-generation scripts, and figure input files are available at https://github.com/pangenome/phrs.

The chromosome-contact and annotation datasets we reanalyzed are public. HPRC v2 Hi-C (CHM13, HG002 and four further samples), CHM13 Arima Hi-C, and the CHM13v2.0 RefSeq/Liftoff v5.2 annotation are in the HPRC data tables and S3 bucket (https://github.com/human-pangenomics/hprc_intermediate_assembly); HG002 Pore-C is the UCSC/GIAB R10.4.1 nanopore Pore-C release in the same bucket. HG002 CiFi is ENA run ERR14656265 in BioProject PRJEB83708 [34]. Mouse meiotic Hi-C is GEO GSE155638 (runs SRR12376885–SRR12376892) [36]. The B6 and CAST T2T assemblies are in ENA BioProject PRJEB47108 (GCA 964188535 and GCA 964188545) [35], and the WashU three-generation T2T pedigree assemblies are at https://github.com/biomonika/washu-pedigree [37].

## Extended Data Figure Legends

**Fig. ED1:**
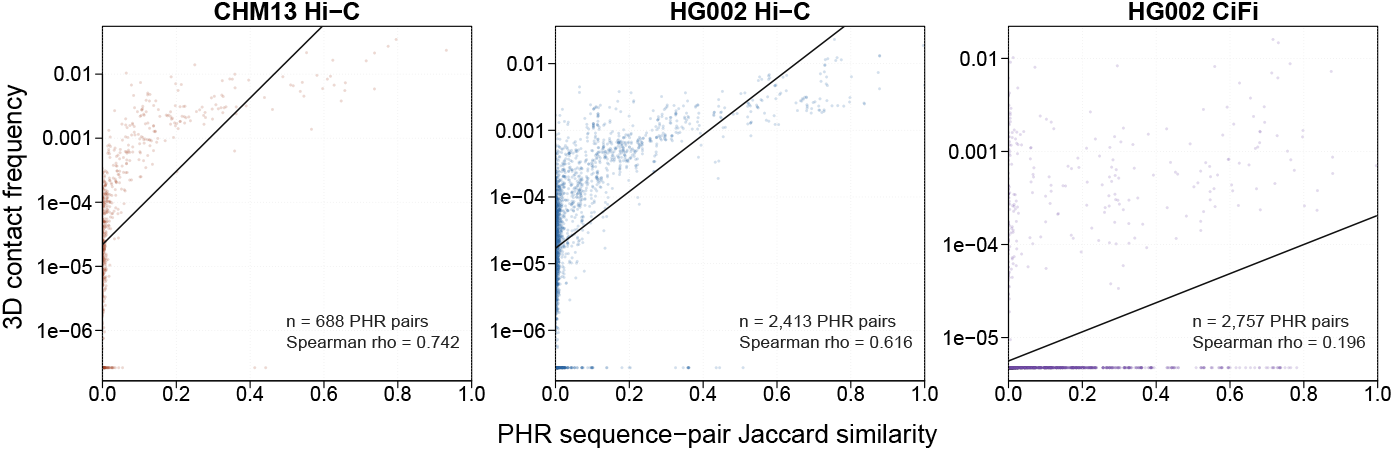
The sequence-to-3D relationship replicates across genomes and contact assays. Three companions to Fig. 4A, each panel a different genome and contact assay (two genomes across three assays), with axes and display as in Fig. 4A. Contact rises with similarity in every panel, reported as pointwise Spearman effect sizes: CHM13 Hi-C (a second genome; *ρ* = 0.742, *n* = 688), HG002 Hi-C (same genome as Fig. 4A, different assay; *ρ* = 0.616, *n* = 2,413), and HG002 CiFi (*ρ* = 0.196, *n* = 2,757; CiFi is the sparsest assay, *∼*90% of pairs zero-contact, consistent with the weakest effect size).

**Fig. ED2:**
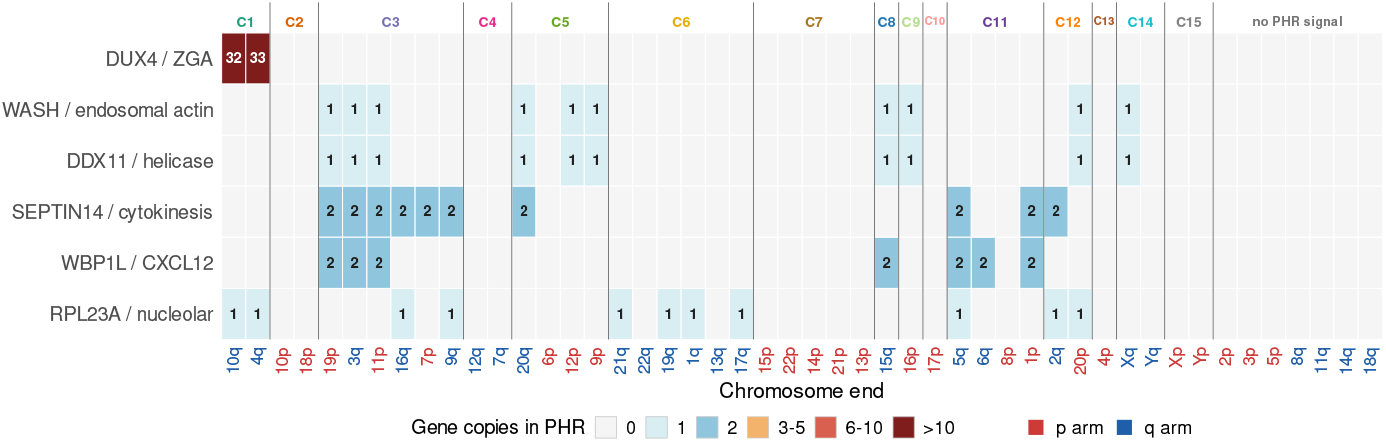
Copy-number-aware functional enrichment across chromosome ends. Heatmap showing the number of CHM13 gene copies in each of six summary functional classes whose midpoints fall within PHRs. Columns are chromosome ends. The 41 PHR-bearing arms are grouped by arm-level PHR community C1–C15 and ordered within each community by the subtelomeric-similarity tree; the seven PHR-free arms are appended at right in gray. Cell values are gene-copy counts: duplicated copies are counted separately rather than collapsed by gene name or family, and the values are not counts of redundant GO/Reactome terms. The six rows group the significantly enriched ontology terms into DUX4/ZGA/transcription/nuclear-envelope/cell-cycle, WASH/endosomal-actin/exocyst, DDX11/helicase/chromosome, SEPTIN14/septin/cytokinesis, WBP1L/CXCL12 signaling and RPL23A/ribosomal/nucleolar classes.

## Human Pangenome Reference Consortium

Derek Albracht^1^, Ivan A. Alexandrov^2^, Jamie Allen^3^, Alawi A. Alsheikh-Ali^4^, Nicolas Altemose^5^, Casey Andrews^6^, Dmitry Antipov^7^, Lucinda Antonacci-Fulton^1^, Mobin Asri^8^, Marcelo Ayllon^9^, Jennifer R. Balacco^10^, Floris P. Barthel^11^, Edward A. Belter Jr^1^, Halle D. Bender^8^, Andrew P. Blair^8^, Davide Bolognini^12^, Katherine E. Bonini^13^, Christina Boucher^14^, Guillaume Bourque^15,16,17^, Silvia Buonaiuto^18^, Shuo Cao^18^, Andrew Carroll^19^, Ann M. Mc Cartney^8^, Monika Cechova^8^, Mark J.P. Chaisson^20^, Pi-Chuan Chang^19^, Xian Chang^8^, Jitender Cheema^3^, Haoyu Cheng^21^, Claudio Ciofi^22^, Hiram Clawson^8^, Sarah Cody^1^, Vincenza Colonna^18^, Holland C. Conwell^23^, Robert Cook-Deegan^24^, Mark Diekhans^8^, Maria Angela Diroma^22^, Daniel Doerr^25,26,27^, Zheng Dong^6^, Danilo Dubocanin^5^, Richard Durbin^28,29^, Jana Ebler^25,30^, Evan E. Eichler^9,31^, Jordan M. Eizenga^8^, Parsa Eskandar^8^, Eddie Ferro^14^, Anna-Sophie Fiston-Lavier^32,33^, Sarah M. Ford^23^, Willard W. Ford^34^, Giulio Formenti^10^, Adam Frankish^3^, Mallory A. Freeberg^3^, Qichen Fu^6^, Stephanie M. Fullerton^35^, Robert S. Fulton^1^, Shenghan Gao^36^, Yan Gao^37^, Gage H. Garcia^9^, Obed A. Garcia^38^, Joshua M.V. Gardner^8^, Shilpa Garg^39^, Erik Garrison^18^, Nani-baa’ A. Garrison^40,41,42^, John E. Garza^1^, Margarita Geleta^43^, Mohammadmersad Ghorbani^44^, Tina A. Graves-Lindsay^1^, Richard E. Green^23^, Carol W. Greider^45^, Cristian Groza^46^, Bida Gu^20^, Andrea Guarracino^11,18^, Melissa Gymrek^47^, Maximilian Haeussler^8^, Leanne Haggerty^3^, Ira M. Hall^48,49^, Nancy F. Hansen^7^, Yue Hao^11^, Mohammad Amiruddin Hashmi^4^, David Haussler^8^, Prajna Hebbar^8^, Peter Heringer^25,26,27^, Glenn Hickey^8^, Todd L. Hillaker^8^, S. Nakib Hossain^3^, Neng Huang^37,50^, Sarah E. Hunt^3^, Toby Hunt^3^, Alexander G. Ioannidis^5,8^, Nafiseh Jafarzadeh^8^, Nivesh Jain^10^, Erich D. Jarvis^10,31^, Maryam Jehangir^11^, Juan Jiang^6^, Eimear E. Kenny^13^, Juhyun Kim^7^, Bonhwang Koo^10^, Sergey Koren^7^, Milinn Kremitzki^1,6^, Charles H. Langley^51^, Ben Langmead^52^, Heather A. Lawson^6^, Daofeng Li^6^, Heng Li^37,50^, Wen-Wei Liao^48,49^, Jiadong Lin^9^, Tianjie Liu^6^, Glennis A. Logsdon^36^, Ryan Lorig-Roach^8^, Jonathan LoTempio Jr^53^, Hailey Loucks^8^, Jane E. Loveland^3^, Jianguo Lu^54^, Shuangjia Lu^48,49^, Julian K. Lucas^8^, Walfred Ma^20^, Juan F. Macias-Velasco^1,6,55^, Kateryna D. Makova^56^, Maximillian G. Marin^37,50^, Christopher Markovic^1^, Tobias Marschall^25,30^, Franco L. Marsico^18^, Fergal J. Martin^3^, Mira Mastoras^8^, Capucine Mayoud^32^, Brandy McNulty^8^, Jack A. Medico^10^, Julian M. Menendez^8^, Karen H. Miga^8^, Anna Minkina^57^, Matthew W. Mitchell^58^, Saswat K. Mohanty^59^, Younes Mokrab^44,60,61^, Jean Monlong^62^, Shabir Moosa^44^, Avelina Moreno-Ochando^63,64^, Shinichi Morishita^65^, Jonathan M. Mudge^3^, Katherine M. Munson^9^, Njagi Mwaniki^66^, Nasna Nassir^4^, Chiara Natali^22^, Shloka Negi^8^, Lingbin Ni^9^, Adam M. Novak^8^, Faith Okamoto^8^, Keisuke K. Oshima^36^, Pilar N. Ossorio^67^, Chie Owa^65^, Sadye Paez^10^, Benedict Paten^8^, Clelia Peano^12,68^, Adam M. Phillippy^7^, Brandon D. Pickett^7^, Laura Pignata^18^, Nadia Pisanti^66^, David Porubsky^9,69^, Pjotr Prins^18^, Timofey Prodanov^25,30^, Anandi Radhakrishnan^8^, T. Rhyker Ranallo-Benavidez^11^, Brian J. Raney^8^, Mikko Rautiainen^70^, Alessandro Raveane^12^, Andreas Rechtsteiner^45^, Luyao Ren^9,31^, Arang Rhie^7^, Fedor Ryabov^71,72^, Samuel Sacco^23^, Farnaz Salehi^18^, Michael C. Schatz^52,73^, Laura B. Scheinfeldt^74^, Aarushi Sehgal^34^, William E. Seligmann^23^, Mahsa Shabani^75^, Kishwar Shafin^19^, Shadi Shahatit^32^, Ruhollah Shemirani^13^, Vikram S. Shivakumar^52^, Swati Sinha^3^, Jouni Siŕen^8^, Linńea Smeds^59^, Steven J. Solar^7^, Marco Sollitto^10,22^, Nicole Soranzo^12,28,76^, Andrew B. Stergachis^9,57^, Marie-Marthe Suner^3^, Yoshihiko Suzuki^65^, Arda Söylev^25,30^, Ahmad Abou Tayoun^77,78^, Jack A.S. Tierney^3^, Chad Tomlinson^1^, Francesca Floriana Tricomi^3^, Mohammed Uddin^4,79^, Matteo Tommaso Ungaro^23,80^, Rahul Varki^14^, Flavia Villani^18^, Ivo Violich^8^, Mitchell R. Vollger^57^, Brian P. Walenz^7^, Charles Wang^81^, Lisa E. Wang^13^, Ting Wang^1,6,55^, Aaron M. Wenger^82^, Conor V. Whelan^10^, Zilan Xin^6^, Zheng Xu^6^, Kai Ye^83^, DongAhn Yoo^9^, Wenjin Zhang^6^, Ying Zhou^37^, Xiaoyu Zhuo^6^, Giulia Zunino^12^

1. McDonnell Genome Institute, Washington University School of Medicine, St. Louis, MO 63108, USA

2. Department of Human Molecular Genetics and Biochemistry, Faculty of Medical and Health Sciences, Tel Aviv University, Tel Aviv 69978, Israel

3. European Molecular Biology Laboratory, European Bioinformatics Institute (EMBL-EBI), Wellcome Genome Campus, Hinxton, Cambridge CB10 1SD, UK

4. Center for Applied and Translational Genomics (CATG), Mohammed Bin Rashid University of Medicine and Health Sciences, Dubai Health, Dubai, UAE

5. Department of Genetics, Stanford University, Palo Alto, CA 94304 USA

6. Department of Genetics, Washington University School of Medicine, St. Louis, MO 63110, USA

7. Genome Informatics Section, Center for Genomics and Data Science Research, National Human Genome Research Institute, National Institutes of Health, Bethesda, MD 20892, USA

8. UC Santa Cruz Genomics Institute, University of California, Santa Cruz, CA 95060, USA

9. Department of Genome Sciences, University of Washington School of Medicine, Seattle, WA 98195, USA

10. The Vertebrate Genome Laboratory, The Rockefeller University, New York, NY 10065, USA

11. Bioinnovation and Genome Sciences, The Translational Genomics Research Institute (TGen), Phoenix, AZ 85004, USA

12. Human Technopole, Milan, Italy

13. Institute for Genomic Health, Icahn School of Medicine at Mount Sinai, New York, NY 10029, USA

14. Department of Computer and Information Science and Engineering, University of Florida, Gainesville, FL 32611, USA

15. Canadian Center for Computational Genomics, McGill University, Montŕeal, QC H3A 0G1, Canada

16. Department of Human Genetics, McGill University, Montŕeal, QC H3A 0G1, Canada

17. Victor Phillip Dahdaleh Institute of Genomic Medicine, Montŕeal, QC H3A 0G1, Canada

18. Department of Genetics, Genomics and Informatics, University of Tennessee Health Science Center, Memphis, TN 38163, USA

19. Google LLC, Mountain View, CA 94043, USA

20. Quantitative and Computational Biology, University of Southern California, Los Angeles, CA 90089, USA

21. Department of Biomedical Informatics and Data Science, Yale School of Medicine, New Haven, CT 06510, USA

22. Department of Biology, University of Florence, Sesto Fiorentino, FI 50019, Italy

23. Department of Ecology and Evolutionary Biology, University of California, Santa Cruz, CA 95060, USA

24. Arizona State University, Consortium for Science, Policy & Outcomes, Washington, DC 20006, USA

25. Center for Digital Medicine, Heinrich Heine University Düsseldorf, Düsseldorf, NRW, DE

26. Department for Endocrinology and Diabetology at the Medical Faculty and University Hospital Düsseldorf, Heinrich Heine University Düsseldorf, Düsseldorf, NRW, DE

27. Paul-Langerhans-Group Computational Diabetology, German Diabetes Center (DDZ) and Leibniz Institute for Diabetes Research, Düsseldorf, NRW, DE

28. Wellcome Sanger Institute, Genome Campus, Hinxton, CB10 1RQ, UK

29. Department of Genetics, University of Cambridge, Cambridge, CB2 3EH, UK

30. Institute for Medical Biometry and Bioinformatics, Medical Faculty and University Hospital Düsseldorf, Heinrich Heine University, Düsseldorf, NRW, DE

31. Howard Hughes Medical Institute, Chevy Chase, MD 20815, USA

32. ISEM, Univ Montpellier, CNRS, IRD, Montpellier, FR

33. Institut Universitaire de France, Paris, FR

34. Department of Computer Science and Engineering, University of California San Diego, La Jolla, CA 92093, USA

35. Department of Bioethics & Humanities, University of Washington School of Medicine, Seattle, WA 98195, USA

36. Department of Genetics, Epigenetics Institute, Perelman School of Medicine, University of Pennsylvania, Philadelphia, PA 19104, USA

37. Department of Data Science, Dana-Farber Cancer Institute, Boston, MA 02215, USA

38. Department of Anthropology, University of Kansas, Lawrence, KS 66045, USA

39. School of Health Sciences, University of Manchester, Manchester M13 9PL, UK

40. Traditional, ancestral and unceded territory of the Gabrielino/Tongva peoples, Institute for Society & Genetics, University of California, Los Angeles, Los Angeles, CA 90095, USA

41. Traditional, ancestral and unceded territory of the Gabrielino/Tongva peoples, Institute for Precision Health, David Geffen School of Medicine, University of California, Los Angeles, Los Angeles, CA 90095, USA

42. Traditional, ancestral and unceded territory of the Gabrielino/Tongva peoples, Division of General Internal Medicine & Health Services Research, David Geffen School of Medicine, University of California, Los Angeles, Los Angeles, CA 90095, USA

43. Department of Electrical Engineering and Computer Science, University of California, Berkeley, Berkeley, CA 94720, USA

44. Medical and Population Genomics Lab, Sidra Medicine, Doha, Qatar

45. Department of Molecular Cell and Developmental Biology, University of California, Santa Cruz, CA, USA

46. Montreal Heart Institute, Montŕeal, QC, Canada

47. Department of Pediatrics, University of California San Diego, La Jolla, CA 92093, USA

48. Center for Genomic Health, Yale University School of Medicine, New Haven, CT 06510, USA

49. Department of Genetics, Yale University School of Medicine, New Haven, CT 06510, USA

50. Department of Biomedical Informatics, Harvard Medical School, Boston, MA 02115, USA

51. Department of Evolution and Ecology and the Center for Population Biology, University of California, One Shields, Davis, CA 95616, USA

52. Department of Computer Science, Johns Hopkins University, Baltimore, MD 21218, USA

53. Department of Pediatrics, Division of Genetics, School of Medicine, University of California, Irvine, CA 92697, USA

54. Sun Yat-sen University, Guangzhou, China

55. Edison Family Center for Genome Sciences & Systems Biology, Washington University School of Medicine, St. Louis, MO 63110, USA

56. Department of Biology and Center for Medical Genomics, Penn State University, University Park, PA 16802, USA

57. Division of Medical Genetics, Department of Medicine, University of Washington School of Medicine, Seattle, WA 98195, USA

58. The Jackson Laboratory for Genomic Medicine, Farmington, CT 06032, USA

59. Department of Biology, Penn State University, University Park, PA 16802, USA

60. Department of Biomedical Science, College of Health Sciences, Qatar University, Doha, Qatar

61. Department of Genetic Medicine, Weill Cornell Medicine-Qatar, Doha, Qatar

62. IRSD - Digestive Health Research Institute, University of Toulouse, INSERM, INRAE, ENVT, UPS, Toulouse, FR

63. MATCH biosystems, S.L., Elche, Spain

64. Universidad Miguel Hernández de Elche, Elche, Spain

65. Department of Computational Biology and Medical Sciences, The University of Tokyo, Kashiwa, Chiba 277-8561, Japan

66. Department of Computer Science, University of Pisa, Pisa, Italy

67. Law School, University of Wisconsin-Madison, Madison, WI 53706, USA

68. Institute of Genetics and Biomedical Research, UoS of Milan, National Research Council, Milan, Italy

69. Genome Biology Unit, European Molecular Biology Laboratory (EMBL), Heidelberg, DE

70. Institute for Molecular Medicine Finland, Helsinki Institute of Life Science, University of Helsinki, Helsinki, Finland

71. The Center for Bio-and Medical Technologies, Moscow, RUS

72. Centre for Biomedical Research and Technology, HSE University, Moscow, RUS

73. Department of Biology, Johns Hopkins University, Baltimore, MD 21218, USA

74. Coriell Institute for Medical Research, Camden, NJ 08103, USA

75. University of Amsterdam, Amsterdam, Netherlands

76. School of Clinical Medicine, University of Cambridge, Cambridge, CB2 0SP, UK

77. Center for Genomic Discovery, Mohammed Bin Rashid University, Dubai Health, UAE

78. Dubai Health Genomic Medicine Center, Dubai Health, UAE

79. GenomeArc Inc, Mississauga, ON, Canada

80. Department of Biology and Biotechnologies “Charles Darwin”, University of Rome “La Sapienza”, Rome 00185, IT

81. Center for Genomics, Loma Linda University School of Medicine, Loma Linda, CA 92350, USA

82. PacBio, Menlo Park, CA 94025, USA

83. The first affiliated hospital of Xi’an Jiaotong University, Xi’an Jiaotong University, Xi’an, Shaanxi, 710049, China

## Funding Statement

We would like to acknowledge the National Human Genome Research Institute (NHGRI) for funding the following grants supporting the creation of the human pangenome reference: U41HG010972, U01HG010971, U01HG013760, U01HG013755, U01HG013748, U01HG013744, R01HG011274, and the Human Pangenome Reference Consortium (BioProject ID: PRJNA730823). This research was supported in part by the Intramural Research Program of the National Institutes of Health (NIH). The contributions of the NIH author(s) are considered Works of the United States Government. The findings and conclusions presented in this paper are those of the author(s) and do not necessarily reflect the views of the NIH or the U.S. Department of Health and Human Services. This work utilized the computational resources of the NIH HPC Biowulf cluster (https://hpc.nih.gov).

## Notes

https://github.com/pangenome/phrs

### Competing Interest Statement

The authors have declared no competing interest.

### Summary of Updates

Addition of gene enrichment analysis, improvement of pedigree study, improvement of manuscript text, discussion, and presentation of results.

